# Selection For Yield Enhanced Rhizobial Mutualism In Pea

**DOI:** 10.64898/2026.05.15.725492

**Authors:** Niall Millar, Clarice Coyne, Stephanie Porter

## Abstract

Crop improvement can enhance food security, but side effects, such as trade-offs between valuable agronomic traits, are common. Likewise, fertilisation helps ensure high yields, but can devalue mutualisms with soil microbes that would otherwise be essential for nutrient acquisition. If the need for nutritional mutualisms is reduced in crops, mutualisms could be disrupted by selection relaxation or allocation trade-offs. Thus, crops could achieve high yields in spite of, or because of, disruption of nutritional mutualisms. Alternatively, the highest-yielding plants might flourish because they maximise nutrient acquisition from both symbionts and the soil. Here, enhanced mutualism could evolve over the course of agricultural crop improvement. To investigate whether high yields in cultivars and wild accessions are negatively or positively genetically correlated with outcomes in the legume-rhizobia mutualism, we measured whether yield and symbiosis traits trade-off or are positively genetically correlated among cultivars and wild accessions. We also tested whether this relationship differs between accessions released before or after 1950. We measured genetic correlations between yield and mutualism traits in 87 domesticated pea (*Pisum sativum)* accessions in a common garden agricultural field across three years. Seed yield and N_2_ fixation (%Ndfa) were positively genetically correlated. While N fixation was more strongly predictive of yield in the pre-1950 accessions than the post-1950 accessions, the underlying positive genetic correlation between the traits did not differ between the groups. The positive genetic correlation between yield and N_2_ fixation indicates that selection to increase yields has maintained or increased the benefits of the rhizobial mutualism in pea. Our findings predict that breeding to increase yield may continue to produce pea cultivars that get a greater proportion of their N from rhizobia, enhancing symbiotic mutualism and reducing the proportion of N supplied by fertilisation.

## Introduction

The improvement of crops for human use is a vital and ongoing effort. Over the 20^th^ century, crop improvement has delivered yearly increases in the yields of staple crops like rice (Peng et al., 2000), wheat (Evans et al., 1980), and legumes (Khazaei et al., 2021). Further gains from breeding techniques are expected to help improve food security for the future (Biswas et al., 2023; Lenaerts et al., 2019). There are, however, side-effects that arise during crop improvement, such as genetic costs due to inbreeding that can increase the frequency of deleterious mutations (Moyers et al., 2018) and trade-offs between traits, such as growth and defence against herbivores (Chen et al., 2015) and pathogens (Córdova-Campos et al., 2012) that can constrain trait evolution. Further, agricultural conditions could allow for the relaxation of selection on traits that could be more critical for performance in the wild than in agriculture (Zohary, 2004). Genetic costs, trade-offs, and relaxed selection are evolutionary factors that could inadvertently disrupt critical plant mutualisms with symbionts such as arbuscular mycorrhizal (AM) fungi (Jacott et al., 2017; Martín-Robles et al., 2018) and nitrogen-fixing rhizobial bacteria (Porter & Sachs, 2020) during crop improvement. Alternatively, if mutualistic symbiosis remains critical to crop performance and yield, enhancement of these mutualisms could contribute to evolutionary processes during crop improvement, either as a direct or indirect target of plant breeding (Porter & Sachs, 2020).

The symbiosis between legumes and rhizobia is critical to biological productivity as it contributes nearly half of all terrestrial N_2_ fixation (Davies-Barnard & Friedlingstein, 2020), a quarter of agronomic crop production, and a third of human-consumed protein (Smýkal et al., 2015). In the rhizobial mutualism, legumes host rhizobia in root nodules, wherein the bacteria fix atmospheric nitrogen (N) into compounds usable by the plant in exchange for carbon (White et al., 2007). Having access to atmospheric N can enhance legume crop productivity (Goyal et al., 2021), and benefit non-legume crops in rotation (Weinert et al., 2023). However, N-enriched conditions in agricultural fields that receive synthetic N-fertilizer could devalue the N provided by rhizobia (Regus et al., 2016), allowing for a relaxation of selection on symbiosis (Porter & Sachs, 2020). Freely available soil-N triggers legumes to reduce investment in symbiotic nitrogen fixation (Nishida & Suzaki, 2018). If reducing investment in symbiotic nitrogen fixation in agricultural conditions enables a legume to allocate more resources to growth and reproduction, this potential resource allocation trade-off could result in a scenario where varieties selected for high yield show low symbiotic function, and manifest as a negative genetic correlation between these traits among varieties. However, the potential for genetic correlations between yield and symbiosis traits have not typically been considered directly in crop breeding decisions (but see (Jacott et al., 2017)).

Trade-offs can shape the evolution of multiple traits in response to natural or artificial selection. Evolutionary trade-offs are measured as a negative genetic correlation between two traits (Roff & Fairbairn, 2007). This could occur if multiple traits are limited by a common resource, resulting in a conflict over its allocation (De Jong & Van Noordwijktt, 1992) or due to antagonistic pleiotropy, whereby a genetic change that increases one trait causes a reduction in another (Mauro & Ghalambor, 2020). During crop improvement, trade-offs during selection for yield can result in reduced chemical defenses (Whitehead & Poveda, 2019) and drought tolerance (Koziol et al., 2012). Similarly, crop improvement may inadvertently reduce crop benefit from mycorrhizal mutualism (Jacott et al., 2017; Martín-Robles et al., 2018; Xing et al., 2012), and control over rhizobial symbionts (Kiers et al., 2007).

Quantifying the strength and direction of genetic correlations between yield traits and symbiosis traits is needed to predict how selection for high yield impacts genetic changes in symbiosis traits. If yield and symbiosis traits trade-off, this may generate negative genetic correlations between yield traits and symbiosis traits among varieties. Here, plant resource allocation to seeds would comes at the expense of allocation to nitrogen-fixing root nodules, and selecting for plants with higher yields could indirectly select for lower symbiotic N_2_ fixation (Tamagno et al., 2018). These improved crops would then get a lower proportion of their N from rhizobia. However, an opposite scenario wherein yield traits and symbiosis traits are positively correlated is also possible. If symbiotic N_2_ fixation instead enhances seed production by boosting total nitrogen acquisition, selecting for plants with higher yields might indirectly select for plants that receive a greater proportion of their N from rhizobia. It is possible to produce pea lines that have both increased yields and N_2_ fixation (%Ndfa) potential, at least when crossing with nodulation mutants (Dhillon et al., 2022). Furthermore, some high yielding domesticated legumes like soybeans can show greater N_2_ fixation than their wild relatives (Muñoz et al., 2016), though they may not receive as much net growth benefit from rhizobia as wild soybeans (Millar et al., 2023). A third possible scenario is that yield and symbiosis traits show no genetic correlation such that breeding to enhance yield does not affect the evolution of rhizobial mutualism. Greater N_2_ fixation does not always result in greater yields (Herridge et al., 2005; Sikinarum et al., 2007), which could occur if the evolution of symbiosis traits is independent and unconstrained by the evolution of yield traits. Understanding which of these hypotheses is best supported will help to inform strategies for enhancement of the rhizobial mutualism in agriculture.

Plant-microbe mutualisms may have unrealised potential as breeding targets (Bennett et al., 2013; Dwivedi et al., 2015). Improvements to the rhizobial mutualism could create more sustainable legume crops that require less N fertilisation for the same or greater yields. For example, crop breeding for N_2_ fixation capacity in soybeans, common beans, and pigeon pea shows promise for increasing yields (Alves et al., 2003; Herridge et al., 1998; Nicolás et al., 2002). Peas have been characterised as poor N fixers in comparison to other legume crops (Liu et al., 2019), so they stand to reap major benefit from similarly targeted breeding programs. Symbiotic mutants of pea can show both improved N_2_ fixation characteristics (Sidorova et al., 2011) and yield (Dhillon et al., 2022). However, mutants with increased nodulation don’t always show greater N_2_ fixation (Voisin et al., 2015), and when they do, it does not necessarily increase yields (Yang et al., 2017). So, while there is potential for the yields and the relatively low N_2_ fixation rates of peas to be improved (Liu *et al*., 2019), these outcomes are not guaranteed to cooccur when generating symbiotic mutants. While there can be a strong phenotypic correlation between yield and N_2_ fixation in pea (Kumar & Goh, 2000), little information addresses the possibility of a genetic correlation that would allow for the improvement of both traits through selection, or cause a trade-off between them.

To uncover the evolutionary trajectory of beneficial symbiosis over the course of domestication and crop improvement in pea, we characterize diverse pea accessions under field conditions to distinguish, 1) whether genetic correlations between yield traits and symbiosis traits are: a) negative, consistent with resource allocation trade-offs or relaxed selection on symbiosis, b) positive, consistent with indirect selection for increased symbiotic function, or c) undetectable, consistent with no impact of N_2_ fixation on yield. To understand how crop improvement affects trait correlations we asked, 2) whether the genetic correlation between yield and symbiosis traits changed over the course of crop improvement during the 20^th^ century. To answer these questions, we used pea accessions from the “More Pea, More Protein, More Profit” (MP3) *Pisum sativum* diversity panel (Cheng et al., 2015; Jermyn & Slinkard, 1977). This panel of 487 pea accessions were selected for high seed protein concentration . We focused our investigation of symbiosis traits on a subset of 87 accessions from this population selected to span the diversity present in the MP3 panel, as well as eleven wild pea genotypes.

## Methods

### Pea germplasm, location, and experiment design

The Pacific Northwest of the USA is a major center of pea agriculture, making it a useful context in which to investigate key crop trait correlations. As part of a broader study to identify pea germplasm useful for breeding peas with high seed protein concentration, 487 MP3 accessions (Cheng et al., 2015; Jermyn & Slinkard, 1977) were planted in a randomised complete block design with three replications, at the USDA Central Ferry Research farm (46°39’5.1’’ N, 117°45’45.4’’ W; 198 m asl) in the Palouse region of eastern WA in 2019, 2020, and 2021 (Sari et al., 2025). We present data for 87 accessions from this broader study of domesticated pea, *Pisum sativum,* that we selected to represent the major groups of pea diversity presented in Holdsworth *et al*. (Holdsworth et al., 2017) and Cheng *et al*. (Cheng et al., 2015) as well as 11 accessions of domesticated pea’s closest wild relative, *Pisum sativum elatius* (Kosterin et al., 2010) (Table S1).

Prior to planting each year, soil samples were taken across the three adjacent spatial blocks in the agricultural field, homogenised, and tested for N, P, K, S, B, Zn, and Mn content (Soiltest Farm Consultants, Moses Lake, WA). Based on deficiencies detected by the tests, fertilisers were applied to the field each year before planting (Table S2). Additionally, preemergence herbicide (α, α, α -trifluoro-2,6-dinitro-*N,N*-dipropyl-P-toluidine; Treflan, Dow Chemical), was applied to the soil prior to planting each year. As is standard for pea cultivation in the region, the seeds were pre-treated with fungicides (mefenoxam [13.3 mL active ingredient (a.i.) 45 kg^−1^], fludioxonil [2.4 mL a.i. 45 kg^−1^], and thiabendazole [82.9 mL a.i. 45 kg^−1^]) insecticide (thiamethoxam [14.3 mL a.i. 45 kg^−1^]), and sodium molybdate (16 g 45 kg^−1^).

The experiment was planted on the 16^th^, 8^th^, and 1^st^ of April across the three years of the experiment, respectively, to optimize weather conditions for planting. In each of the three years of the experiment, there were three adjacent blocks containing 487 plots each, one for each accession, for a total of 1461 plots per year. We focused on a subset of 87 accessions of domesticated pea from this larger experiment (Table S1), so each year we sampled from 87 randomly placed plots per block, for a total of 261 plots per year. In 2020, there were twelve additional plots per block for the eleven wild *P. sativum elatius* accessions and a non-nodulating *P. sativum* accession called Frisson p56 (Sagan *et al*., 1993, Table S1), for a total of 297 plots in that year.

Each plot was a 152 by 30 cm space sown with 30 seeds. The seeds were mechanically drilled in two 152 cm lines with 30 cm center spacing. Plots in the same row were 152 cm apart, and adjacent rows were 100 cm apart. Each row contained 30 plots (excluding the final row in each block, which had fewer than 30 plots), and there were 17 rows per block, for a total of 51 rows per year. Each row was irrigated 10 minutes per day from drip tape buried 15 cm under the center of each plot. On average, 24.4 plants germinated and survived per plot. Peas have been grown at this site for at least 25 years. The soil contains rhizobia compatible with peas so no rhizobial inoculation was applied in this experiment.

In summers (April through August) of all three years, the average monthly rainfall was 1.19 cm. However, in May, 2020, the experiment experienced an extreme weather event when a storm deposited 7.79 cm of rainfall, mostly between the 17^th^ and 20^th^ of the month, giving May 2020 the highest rainfall in the three summers of the study. Storm damage to each plot was assigned a damage severity value (1: No damage – 5: severe damage), which was strongly correlated with yield (Pearson correlation coefficient = -0.64, *P*=0, S1).

### Yield traits

To measure peas’ yield traits, we took measurements aggregated across all individuals in an experimental plot for all three years of the study. Each year the plots containing mature pea seeds were manually harvested in order of maturity, threshed, and the seeds blown clean Generally, this took place between the beginning of the second week of July, and the end of the second week of August. Yield was measured as the total weight (g) of threshed seeds from a plot. Seeds from a plot were ground to powder, homogenized, and analysed for seed protein concentration (SPC) with near-infrared (NIR) spectrometry (Matrix I, Bruker Co.).

In addition to these plot-aggregated phenotypes, we also sampled one individual plant per plot in 2020. In order to measure yield traits for individual plants, on the 19^th^ and 20^th^ of May, 2020, 6 weeks after planting, we sampled a single plant from each replicate plot for 87 accessions, the 11 wild accessions, and the non-nodulating mutant. From each plot, where possible, we chose a plant that was spatially separate from the others, to minimise damage to adjacent plants. We carefully excavated the plant, along with its root system, washed the roots with water, and separated roots from shoots. We wrapped individual root systems in wet paper towels and transported them in chilled coolers before counting the root nodules. We then dried and measured the shoot and root weight of each plant. These individual plants were harvested at the start of the storm in May of 2020, before storm damage occurred.

### Symbiosis traits

Our primary measure of symbiotic nitrogen fixation was the percent of nitrogen derived from the atmosphere (%Ndfa) in pea tissue, which we calculated using the ^15^N natural abundance method (Unkovich et al., 2008),

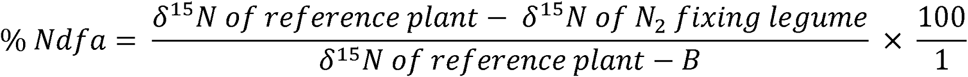

Here,δ^15^, refers toδ^15^N for a pea plant that is entirely dependent on soil N, and *B* is the δ^15^N measured for a pea plant that is entirely dependent on symbiotic N_2_ fixation.

1-1.5 mg samples of the ground, homogenized total seed yield from each plot were encapsulated for stable isotope analysis at the WSU Stable Isotope Core Laboratory in Pullman, WA. They were analysed for δ^15^N, % N, δ^13^C, and % C, by weight. The samples were converted to N_2_ and CO_2_ with an elemental analyser (ECS 4010, Costech Analytical, Valencia, CA); these two gases are separated with a 3m GC column and analysed with a continuous flow isotope ratio mass spectrometer (Delta PlusXP, Thermofinnigan, Bremen). Using the % by weight values for protein, N, and C, we calculated the absolute weight (g) of each of these properties, based on total seed weight. We used the plot-aggregated seed δ^13^C values as estimates of the plants’ water use efficiency (Kaler et al., 2018).

Initial calculations of %Ndfa using the mean δ^15^N value of the non-nodulating mutant as the reference value, and the peas’ lowest δ^15^N value (-2) as the B value, returned some %Ndfa values >100, and <0. In order to obtain %Ndfa values between 0 and 100, we used the δ^15^N value of barley (8.9) (Unkovich et al., 2008), and a B value of -4.5. Therefore, we primarily interpret our %Ndfa values to show relative differences between our samples rather than precise absolute values, since different constants will change the magnitude, but not the ranking of %Ndfa values (Cox et al., 2022). We measured the plot-aggregated %Ndfa of the ground seeds from each plot for each of the three years of the study. In 2020, we also estimated %Ndfa, as above, with a 1-1.5 mg sample of tissue from the apical leaf from the single plant we collected from each plot. We also counted root nodules for this plant.

### Genetic Correlations

The complete randomized block design of this experiment allowed us to partition total phenotypic variance (*V_p_)* into genetic variance components (*V_G_*) and environmental variance components (*V_E_*) (Hill & Mackay, 2004). We thus estimated the proportion of variation in the peas’ yield traits and their symbiosis traits attributable to: 1) *V_G_* as broad sense heritable genetic variation among accessions and, 2) *V_E_* as trait plasticity of accessions due to both spatial environmental variation among blocks in the field (eg due heterogeneity in soil chemistry (Table S2)), and temporal environmental variation among the years of the experiment (e.g. due to weather conditions that year; Fig. S1) .

To calculate genetic correlations among plot-aggregated phenotypes, we applied the following generalized linear mixed model (GLMM) to each of our measured traits (Skovbjerg et al., 2020), which we will call “Model A”:

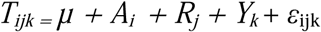

Where *T_ijk_* is a variable representing a given trait, *µ* is the overall mean*, A_i_* is the random effect of the *i*th accession (87 levels), *R_j_* is the random effect of the *j*th experimental block (3 levels per year), *Y_k_* is the fixed effect of experiment year (3 levels), and _εijk_ is the residual error term of the *i*th accession in the *j*th block in the *k*th year. Due to extensive storm damage in 2020, we excluded data from this year in models of genetic correlations with seed %Ndfa, keeping 2019 and 2021. To estimate the best linear predictors (BLUPs) for each trait, we extracted the random effects estimates for accession from each model (Skovbjerg et al., 2020).

Pea varieties are self-fertile inbred lineages, so correlations among accessions’ traits can be viewed as “broad sense genetic correlations” sensu Shaw (1986), as additive genetic effects remain confounded with maternal genetic and epistatic variance. Therefore, the genetic effects we estimate represent an upper bound on the additive genetic variance. To measure broad sense genetic correlations we generated Pearson correlation coefficients between the yield trait BLUPs and the symbiosis trait BLUPs. To account for multiple tests for genetic correlations between yield traits and symbiosis, we subjected all *P*-values for genetic correlations to a sequential Holm-Bonferroni correction.

To test whether yield-symbiosis correlations changed over the course of crop improvement during the 20^th^ century, we separated the accessions into age groups that were introduced before or after 1950. We then generated separate pre- and post-1950 BLUPs, using Model A, but we ran it separately for pre- and post-1950 age groups. As above, we calculated Pearson coefficients for genetic correlations between yield traits and symbiosis traits. We used the Cocor package (version 1.1-4, Diedenhofen & Musch, 2015) to test whether genetic correlations differed between the age groups. Cocor uses Fisher’s *z* to test whether there are differences between correlation coefficients, and whether the difference between the R-squared values exceeds Zou’s confidence intervals (Zou, 2007). Fisher’s *z* and Zou’s confidence intervals test for differences in correlation strength, i.e. the distance of points from the line of best fit. To test for differences in the slopes of the pre- and post-1950 yield-symbiosis correlations, we used a GLMM with the seed %Ndfa BLUPs as the response variable, the fixed effect of the yield BLUPs, age group, their interaction, and the random effect of experimental year. We used a likelihood ratio test to determine the significance of the yield BLUPs: age group interaction, from which we inferred whether the slope of the yield-symbiosis correlation differed between age groups. To explore how influential a set of particularly low values were for the overall model, we conducted a post-hoc test where we removed the five lowest yielding pre-1950 accessions and re-ran the pre/post-1950 correlation comparisons.

In addition to comparing yield-symbiosis correlations between the age groups, we tested the effect of age group on individual traits to see if, for example, seed yield means differ significantly between pre- and post-1950 accessions. We used the following GLMM for each trait (Skovbjerg et al., 2020), which we will call “Model B”:

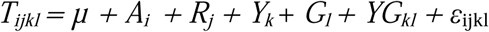

Where *T_ijk_* is a variable representing a given trait, *A_i_* is the random effect of the *i*th accession (87 levels), *R_j_* is the random effect of the *j*th experimental block (3 levels per year), *Y_k_*is the fixed effect of the *k*th experiment year (3 levels), *G_l_*is the fixed effect of an accession’s age group (pre- or post-1950), Y*G_kl_*is the interaction between age group and experimental year, and _εijk_ is the residual error term of the *i*th accession in the *j*th block in the *k*th year, and *µ* is the overall mean. We used likelihood ratio tests to generate χ^2^ values and determine whether trait means were significantly affected by age group.

Traits that were only measured on individual plants in 2020 were modelled separately. For the 2020 single-plant data (including shoot weight, root weight, nodule count, leaf %Ndfa, leaf % by weight N, and total N content), we used Model A for generating BLUPs, and Model B to test the significance of age group on traits. However, we removed the experimental year effect for the 2020-only models.

All analyses were conducted in R version 4.2.2 (R Core Team, 2014). We used the glmmTMB package (version 1.1.14, Brooks et al., 2017) to run GLMMs. We used the DHARMa package (version 0.4.6, Hartig, 2022) to check model assumptions.

## Results

### Genetic correlations between yield and symbiosis traits

In 2019 and 2021 there were positive genetic correlations between seed %Ndfa and the following plot-aggregated traits: yield (r(85) = 0.56, *P*< 0.001, Table S3, Fig. 1a), seed protein weight (r(85) = 0.52, *P*< 0.001, Table S3, Fig. 1b), seed N weight (r(85) = 0.61, *P*< 0.001, Table S3, Fig. 1c), and seed C weight (r(85) = 0.75, *P*< 0.001, Table S3, Fig. 1d). However, seed %Ndfa did not correlate with seed protein concentration, % by weight N, or % by weight C, nor with seed δ^13^C. The total weight of N derived from symbiotic fixation (g Ndfa), calculated from seed %Ndfa and total seed weight (yield), was positively correlated with yield (r(85) = 0.94, *P*= 0, Table S3, Fig. 2a), seed protein weight (r(85) = 0.89, *P*= 0, Fig. 2b), seed N weight (r(85) = 0.99, p=0, Fig. 2c), and seed C weight (r(85) = 0.85, *P*= 0, Fig. 2d). g Ndfa was uncorrelated with seed protein concentration, % by weight N, % by weight C, or seed δ^13^C.

**Fig. 1:**
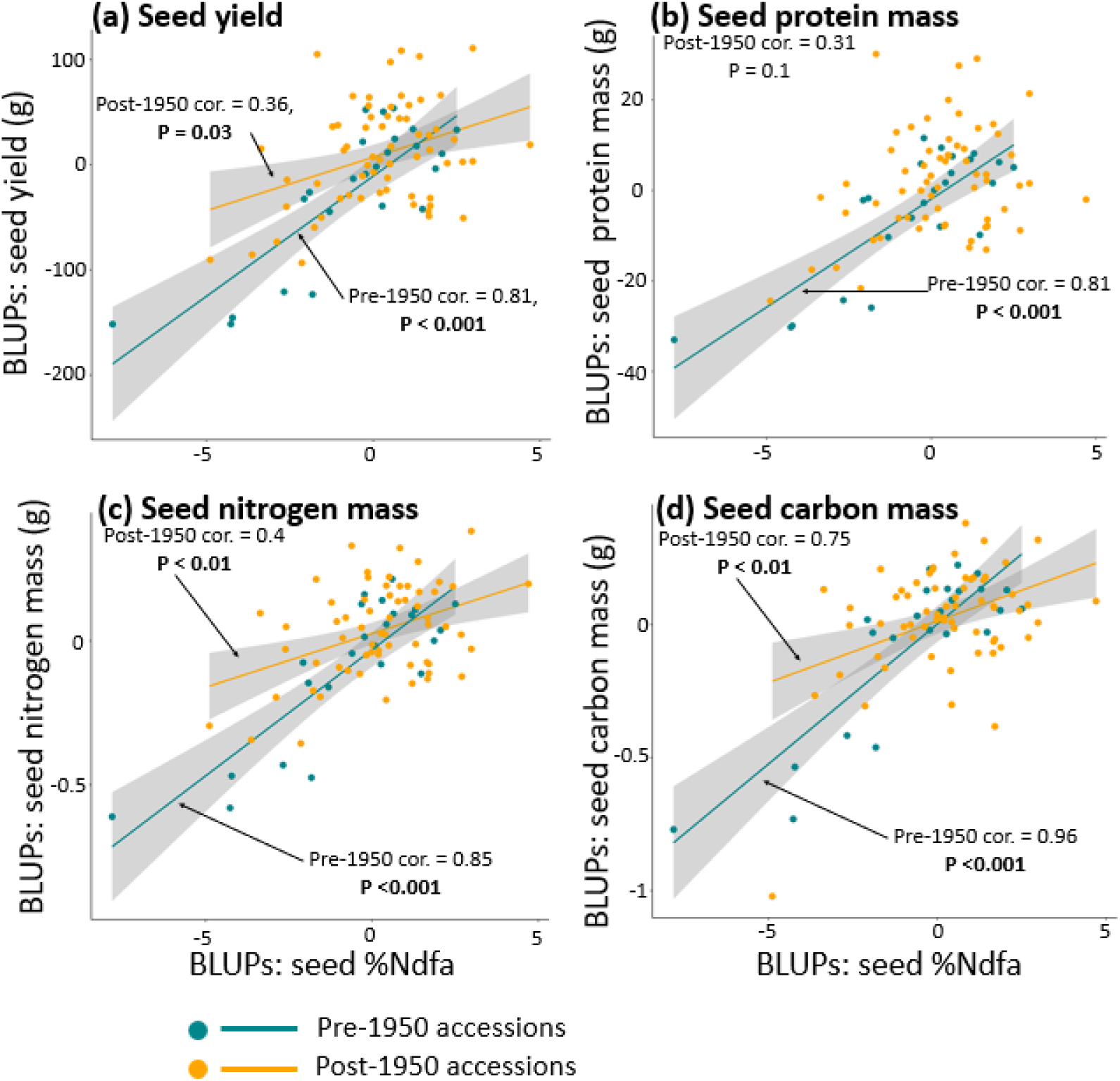
Seed yield is positively genetically correlated with seed %Ndfa. Shown are the best linear unbiased predictors (BLUPs) for each of 87 pea accessions for respective traits (a-d) measured at the level of field plot from 2019 and 2021. Positive genetic correlations were found between seed %Ndfa and (a) seed yield, (b) total seed protein weight, (c) total seed nitrogen weight, and (d) total seed carbon weight, for pre-1950 accessions (blue) and post-1950 accessions (orange). The Pearson correlation coefficients and *P*-values of each pre- and post- 1950 correlation are shown. For all of the correlations shown in this figure, there were significant differences in the strength of the correlation between the pre- and post-1950 accessions.

**Fig. 2:**
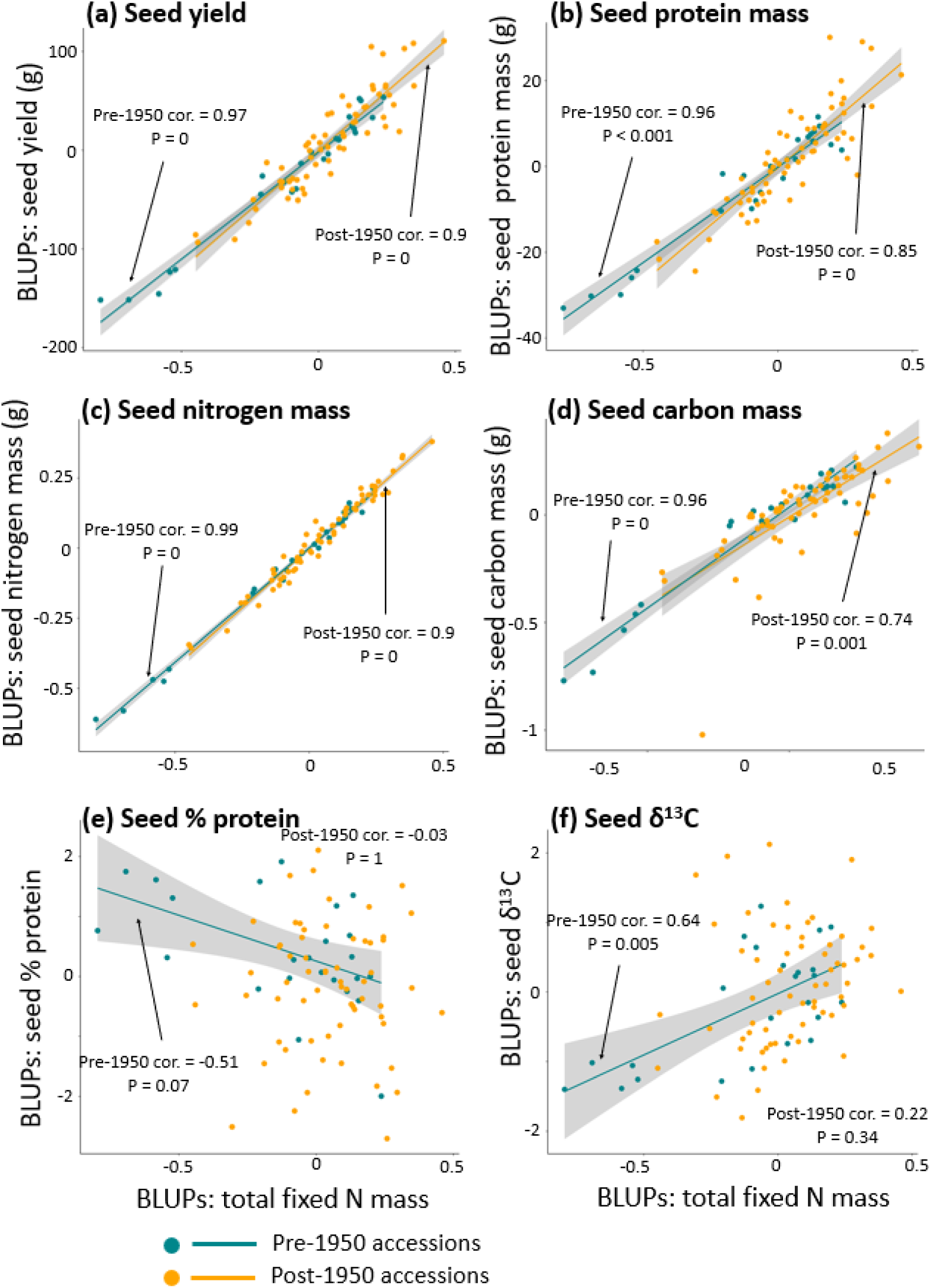
Seed yield is positively genetically correlated with seed g Ndfa. Shown are the best linear unbiased predictors (BLUPs) for each of 87 pea accessions for respective traits (a-f) measured at the level of plot from 2019 and 2021. Positive genetic correlations were found in seed tissues between yield and g Ndfa, δ^13^C, total protein weight, total nitrogen weight, and total carbon weight. There was a marginally significant negative genetic correlation between seed g Ndfa and protein concentration for the pre-1950 accessions. The Pearson correlation coefficients and *P*-values for pre-1950 (blue dots) and post-1950 (orange dots) accessions are shown. There were significant differences between the pre- and post-1950 correlations for all traits.

Due to storm damage in 2020, we do not report correlations between seed %Ndfa and other plot-aggregated traits for this year. However, our single-plant measurements of yield and symbiosis traits were measured on an individual six-week-old plant collected from each plot on May 20, 2020, prior to storm damage. For these individual plant traits, there was a positive genetic correlation between leaf %Ndfa and shoot weight (r(85) = 0.64, *P*<0.001, Table S4, Fig. 3a), and between leaf %Ndfa and shoot N weight (r(85) = 0.65, *P*<0.001, Table S4, Fig. 3b) for domesticated pea. Leaf %Ndfa did not correlate with shoot % by weight N, root weight, or nodule count for domesticated pea. Among the wild accessions grown in 2020, we did not detect a genetic correlation between leaf %Ndfa and shoot weight (Table S7, Fig 3a), nor between leaf %Ndfa and other phenotypes measured on individual plants.

**Fig. 3:**
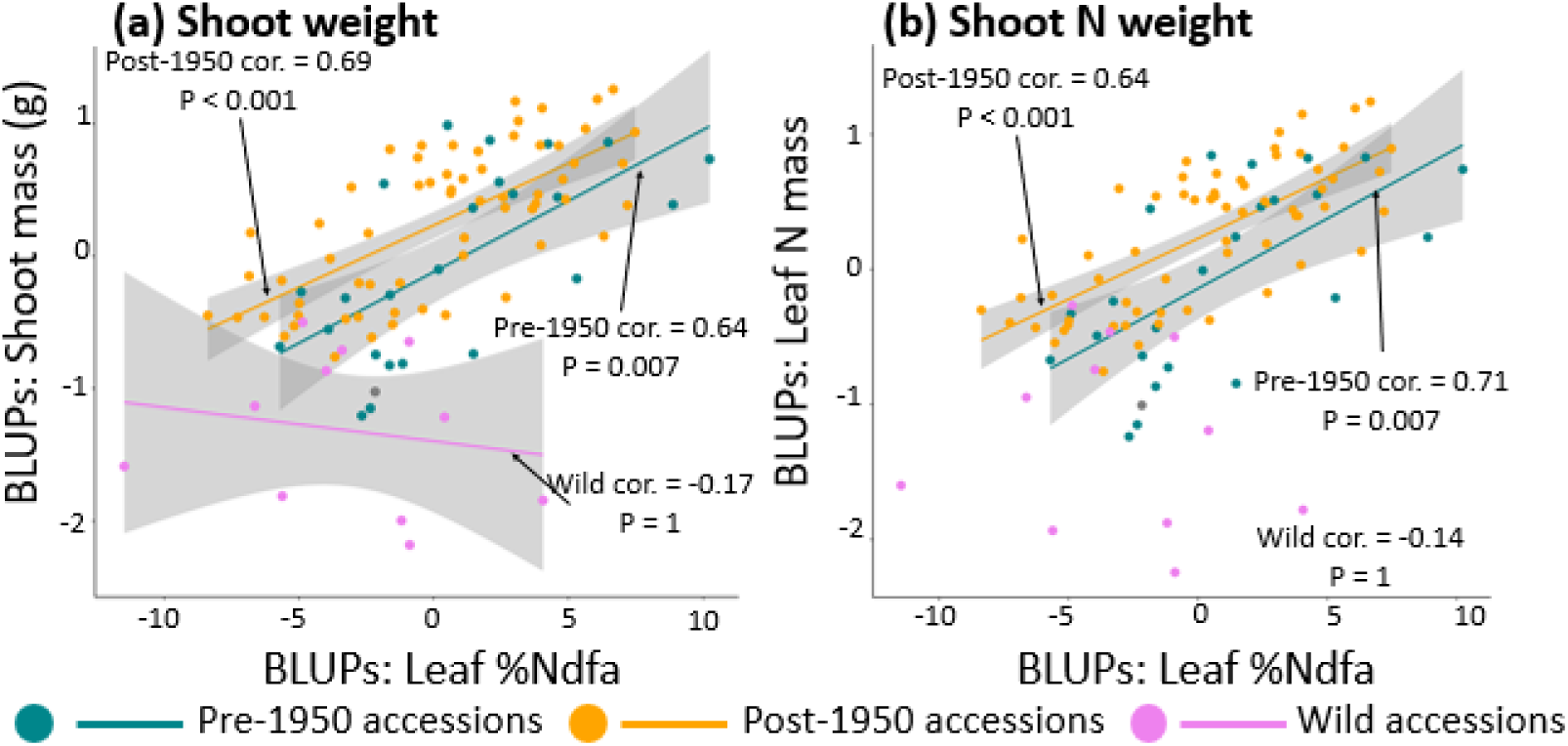
Leaf %Ndfa is positively genetically correlated with shoot weight in domesticated, but not wild peas. Shown are the best linear unbiased predictors (BLUPs) for each of 87 domesticated pea accessions and 11 wild pea accessions for traits measured at the individual plant level in 2020. Positive genetic correlations were found between leaf %Ndfa and shoot weight (a) and leaf %Ndfa and leaf N weight (b) for pre-1950 (blue) and post-1950 (orange) accessions, while leaf %Ndfa and shoot weight were negatively genetically correlated in the wild accessions (violet). Pearson correlation coefficients and *P*-values are shown.

### Genetic correlations in pre- and post- 1950 age groups

To determine whether yield-symbiosis correlations changed over time, we compared separate genetic correlations in accessions registered with the USDA either before or after 1950. For the plot-aggregated data, in both pre- and post-1950 age groups there was a positive correlation between seed yield and seed %Ndfa, although the pre-1950 correlation coefficient (pre-150 r(22)=0.81, *P*<0.001, Table S5a, Fig. 1) was greater than the post-1950 correlation coefficient (post-1950 r(61)=0.36, *P*=0.03, Table S5a, Fig. 1). The difference between the pre- and post-1950 correlations was significant (Fisher’s *z* = 3.03, *P*=0.002, Fig. 1, Table S5a). Similarly, there were stronger correlations in the pre-1950 age group than the post-1950 age group between seed %Ndfa and seed protein weight (Fisher’s *z* = 3.26, *P*=0.001, Fig. 2, Table S5a), N weight (Fisher’s *z* = 3.28, *P*=0.001, Fig. 1, Table S5a) and C weight (Fisher’s *z* = 3.39, *P*<0.001, Fig. 1, Table S5a). However, with the lowest-yielding pre-1950 accessions removed, there were no difference between the yield-%Ndfa correlations for pre- and post-1950 age groups (Table S5b). Further, the GLMM testing the effect of the yield BLUPs: age group interaction on the seed %Ndfa BLUPs found no difference in the slopes of the pre- and post- 1950 yield-%Ndfa correlations (Table S8). The pre- and post-1950 age groups showed no detectable differences in how their seed %Ndfa correlated with their seed δ^13^C, protein concentration, % N weight, or % C weight. In the single-plant measurements from 2020, there were no differences in correlations between the pre- and post-1950 age groups (Table S6).

### Effect of age group on yield and symbiosis traits

To better understand how yield-symbiosis trait correlations have changed over time, we used Model B to test the effect of age group on each trait, while accounting for the effect of accession, year (excluding 2020), and block. There was a marginal increase in seed %Ndfa in the post-1950 accessions (_χ_^2^: 3.23, *P*=0.07, Fig. 4a, Table S3). All of the weight-based traits had higher mean values in the post-1950 age group than in the pre-1950 age group; these included seed yield (χ ^2^: 3.23, *P*=0.009, Fig. 4b, Table S3), g Ndfa (χ ^2^: 6.48, *P*=0.01, Fig. 4c, Table S3), protein weight (_χ_^2^: 5.05, *P*=0.02, Fig. 4d, Table S3), C weight (C weight (χ ^2^:5.39, *P*=0.02, Fig. 4e, Table S3), and, marginally, N weight (χ ^2^: 3.35, *P*=0.06, Fig. 4f, Table S3). Seed δ^13^C was also greater in the post-1950 age group than the pre-1950 age group (χ ^2^: 8.05, *P*=0.004, Fig. 4, Table S3), which could indicate higher water use efficiency in the newer accessions than the older ones. Despite the increase in total seed protein weight, there was a decrease in seed protein concentration in the post-1950 age group (χ ^2^: 5.41, *P*=0.01, Fig. 4h, Table S3).

**Fig. 4:**
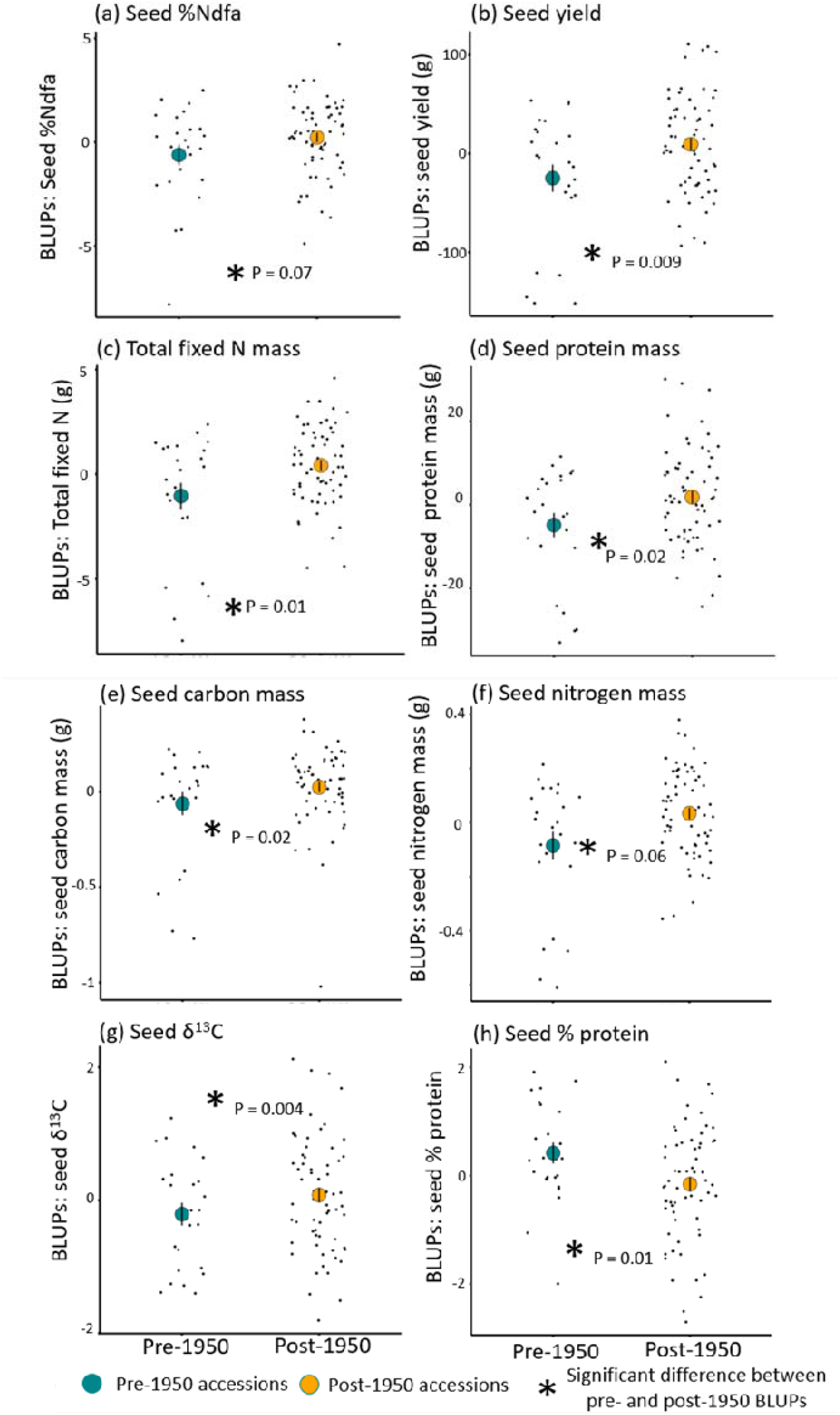
Seed yield measures are greater in post-1950 accessions, while seed protein concentration are greater in pre-1950 accessions. Shown are the best linear unbiased predictors (BLUPs) for each of 87 pea accessions for respective traits (a-d) measured at the level of plot from 2019 and 2021. Seed yield (b), protein weight (d), and carbon weight (e), nitrogen weight (f) were greater in post-1950 accessions than pre-1950 accessions. Seed %Ndfa and g Ndfa were marginally greater in the post-1950 accessions than in post-1950 accessions. Seed protein concentration was greater in pre-1950 accessions than in post-1950 accessions. Significant differences between mean values of age groups are indicated with an asterisk, along with the *P*-value for the effect of age group on that trait. Error bars indicate the standard error of the mean.

For the single-plant measurements in 2020, we compared wild pea accessions to pre- and post-1950 age groups of domesticated pea. Most phenotypes had greater mean values in the domesticated peas, including leaf %Ndfa (χ^2^: 7.95, *P*=0.04, Fig. 5a, Table S7), shoot weight (χ ^2^: 53.7, *P*<0.001, Fig. 5b, Table S7), shoot fixed N weight (χ ^2^: 46.1, *P*<0.001, Fig. 5c, Table S7), and shoot N weight (χ ^2^: 51.8, *P*<0.001, Fig. 5d, Table S7) (seed protein concentration was not measured in the wild accessions). Although shoot N weight was greater in the domesticated accessions, leaf % N was greater in the wild peas (χ ^2^: 8.83, *P*=0.03, Fig. 5e, Table S7). Domesticated root weight was only greater than wild root weight for the post-1950 age group _χ_ : 53.7, *P*=0.01, Fig. 5f, Table S7). Nodule count was indistinguishable among the groups of accessions.

**Fig. 5:**
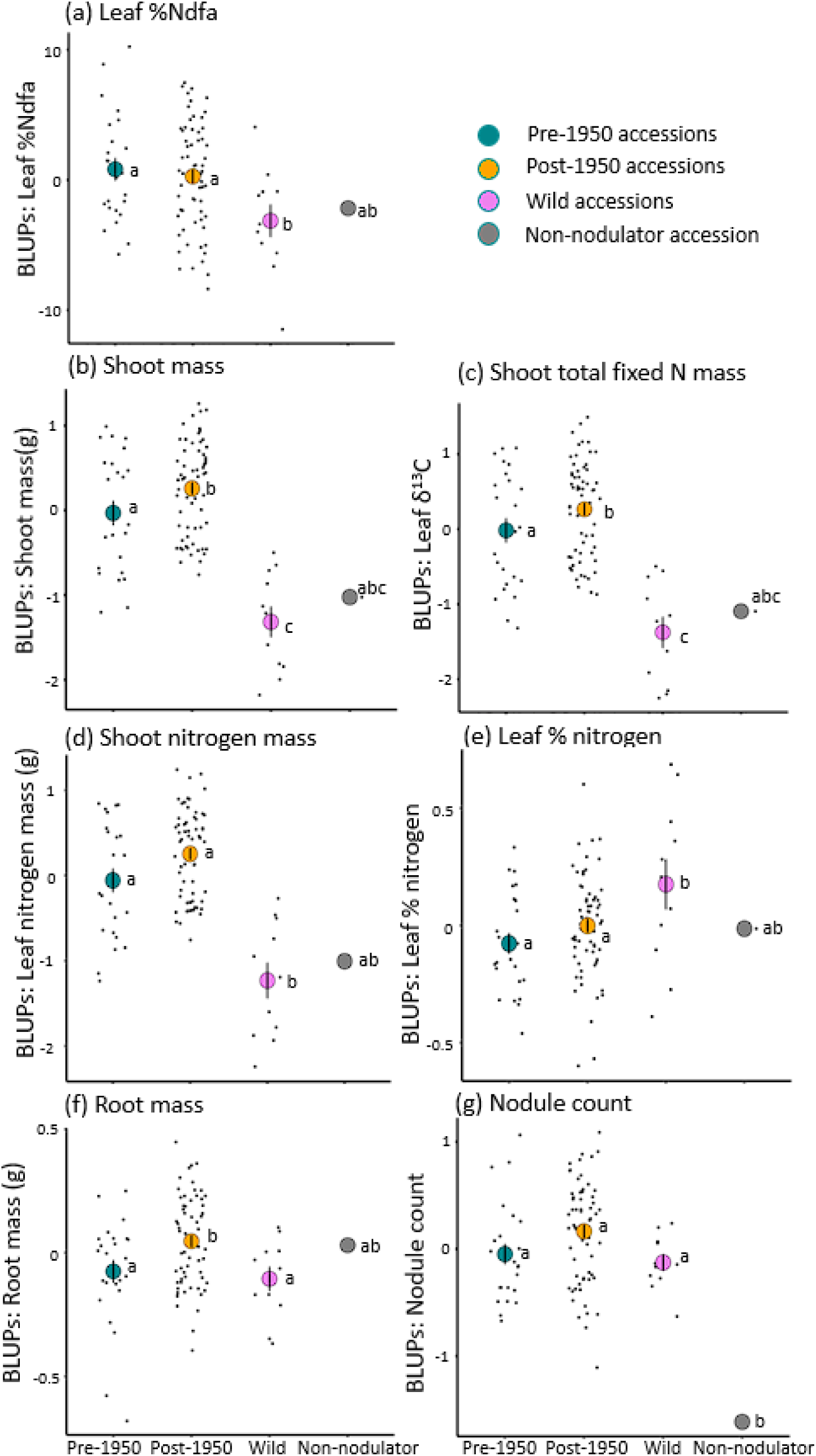
Yield is higher in post-1950 than pre-1950 accessions, and lower in wild peas than domesticated peas. Shown are the best linear unbiased predictors (BLUPs) for each of 87 domesticated pea accessions and 11 wild pea accessions for traits measured at the individual plant level in 2020. Accession groups include those from pre-1950 (blue), post-1950 (orange), the wild (violet), and the non-nodulating accession (grey). Domesticated accessions had higher mean values in the than the wild accessions for most traits, excluding nodule count, which did not differ significantly, and leaf % nitrogen content, which was greater in the wild accessions. Tukey letters indicate pairwise contrasts between the mean values of the age groups. Error bars indicate the standard error of the mean.

Using Model B (excluding the experimental year effect), we tested the effect of pre- or post-1950 age group on the single-plant phenotypes from 2020. Age group had similar effects on the single-plant measurements that were seen in the plot-aggregated measurements from 2019 and 2021: shoot weight (χ ^2^: 4.06, *P*=0.04, Fig. 5b, Table S4), shoot N weight (χ ^2^: 5.18, *P*<0.001, Fig. 5d, Table S4), and root weight (χ ^2^: 6.74, *P*=0.009, Fig. 5f, Table S4) were greater in the post-1950 age group than the pre-1950 age group.

## Discussion

We find that higher-yielding pea accessions obtain a greater proportion of their N from symbiotic N_2_ fixation rhizobia than do lower-yielding accessions. This positive genetic correlation between N_2_ fixation (%Ndfa) and yield in pea supports a scenario in which the evolution of higher yields has enhanced, rather than disrupted mutualistic symbiosis with rhizobia. We infer that artificial selection for increased pea yields has improved the ability of these peas to fix N with rhizobia, rather than a scenario in which an allocation trade-off results in yield gains coming at a cost to legume investment in rhizobial symbiosis. Additionally, the yield increase in pea, and its positive correlation with N_2_ fixation (%Ndfa), appears to have avoided the trade-off with protein experienced by other legumes (Skovbjerg et al., 2020). Therefore, it appears feasible for artificial selection for high yield in pea to continue to generate accessions that enhance both yield and symbiotic N_2_ fixation. However, our findings also reveal abundant standing genetic variation in symbiotic N_2_ fixation. It will thus be critical to avoid inadvertently selecting high yielding lineages with low symbiotic N_2_ fixation for future crop improvement efforts.

Our findings provide a valuable *genetic* perspective of more direct relevance to crop improvement than prior observations of prior positive *phenotypic* correlations between yield and N_2_ fixation as %Ndfa (Dhillon *et al*. 2022) or total fixed N (Dhillon et al., 2022; Kumar & Goh, 2000). This is because heritable, genetically based trait correlations are required to infer evolutionary dynamics. We predict that selection for greater protein weight should not cause an evolutionary trade-off with N_2_ fixation, and could even increase N_2_ fixation, based on the positive genetic correlation between N_2_ fixation and plot-aggregated total seed protein weight. We did not, however, find a genetic correlation between seed protein concentration and N_2_ fixation. It is possible that the genetic correlation between protein weight and N_2_ fixation could arise from the genetic correlation between overall yield and N_2_ fixation. A higher seed protein concentration is a primary goal in pea breeding (Coyne et al., 2005) to facilitate the use of pea protein in food production (Burger & Zhang, 2019; Lu et al., 2020). For these purposes, cultivars that invest a greater proportion of resources in protein production than in overall seed size would be valuable. However, among the MP3 panel of pea diversity we examine, accessions with greater N_2_ fixation do not invest a greater proportion of their resources in protein production, as shown by the lack of a genetic correlation between the two. While other sets of pea diversity that were generated by deliberate crosses with nodulation mutants show positive correlations between N_2_ fixation and seed protein concentration (Dhillon et al., 2022), in the diversity of a panel of conventional pea lines we examine, we do not detect a genetic correlation between greater N_2_ fixation and seed protein concentration, or a trade-off between N_2_ fixation and seed protein concentration. This is promising for pea germplasm improvement compared to other crops like grains where trade-offs with protein are common, such as the trade-off between yield and % protein content in faba bean (Peltonen-Sainio et al., 2011; Curti et al., 2018; Skovbjerg et al., 2019) and between oil and protein in rapeseed (Peltonen-Sainio et al., 2011).

The positive genetic correlation between yield and N_2_ fixation (%Ndfa) and in peas demonstrates that evolution under artificial selection is not necessarily constrained by trade-offs between plant investment in yield and symbiosis. Models often assume that the evolution of mutualisms would be influenced by such trade-offs, such that symbionts like rhizobia could increase their own reproduction by exploiting their host for carbon without fixing N (West et al., 2002)(Porter & Sachs, 2020). The benefits of investing in the symbiosis appear to have been maintained for peas over the course of crop improvement. While these are encouraging results for breeding efforts that seek to improve both agronomic and symbiotic traits in crops (Bennett et al., 2013; Dwivedi et al., 2015), the lack of genetic correlations in the data from 2020 during a major storm, show that the genetic correlation between yield and the rhizobial symbiosis can break down.

Results from 2020 show that the simultaneous evolution of higher yields and greater N_2_ fixation (%Ndfa) evident in 2019 and 2021 could be vulnerable to disruption by extreme weather events. In 2020, an extreme storm event disrupted genetic correlations between yield and symbiosis. In 2020, there was negligible genetic variation in N_2_ fixation (%Ndfa), and no correlation with yield, likely because the storm damage resulted in heterogeneous environmental, rather than genetic, variance for plant traits. Damage from May 2020’s rainstorm was directly predictive of yields: at the end of the growing season the most storm-damaged plots had the lowest yield. Interestingly, the most storm-damaged plots also had the highest N_2_ fixation (%Ndfa). Increases in nodule biomass in response to abiotic stress can occur (Wurzburger & Ford Miniat, 2014). Alternatively, it is possible that pea plants with higher N_2_-fixation (%Ndfa)rates were larger and therefore more susceptible to damage from heavy rain. Correspondingly, individual plants showed a positive correlation between leaf N_2_ fixation (%Ndfa) and shoot weight prior to the storm damage. Weather extremes, and climate change in general, are pervasive threats to agriculture (Aydinalp et al., 2008; Lesk et al., 2016; Vogel et al., 2019). Climate change is a particular concern for plant breeding, since climatic variation can reduce the efficacy of selection and genetic gains made by breeding programs (Xiong et al., 2022). Years such as 2020, with negligible genetic variation in N_2_ fixation (%Ndfa) and no correlation with yield, could interfere with efforts to improve symbiotic benefits in peas.

We observe evolutionary shifts in the genetic correlation between yield and N_2_ fixation (%Ndfa) in pea over the course of the 20^th^ century. The 20^th^ century was a period of intense crop improvement (Bradshaw, 2017) with large increases in yields after 1950, coinciding with the “Green Revolution” and modern agricultural practices (Brancourt-Hulmel et al., 2003; Jin et al., 2010). The 1950s saw the proliferation of artificial fertiliser use and the formation of organisations like the National Pea Improvement Association, so we sought to examine the results of breeding before and after these developments. We find that while both older and newer accessions showed positive genetic correlations between yield and symbiosis, this correlation is stronger in the pre-1950 age group. Taken at face value, this could be a concerning result; post- 1950 breeding efforts seem to have caused the evolution of pea accessions where yield is less closely tied to levels of N_2_ fixation (%Ndfa) than in their predecessors. Such a decoupling of yield and symbiotic resource exchange has been seen for the mycorrhizal symbiosis over the course of crop improvement in wheat (Hetrick et al., 1993; Zhu et al., 2001). However, although yield and N_2_ fixation (%Ndfa) were more closely genetically correlated in the pre-1950 age group than the post-1950 group, the slopes of these correlations do not differ significantly. Therefore, nitrogen fixation rates are less predictive of yield in post-1950 peas, but the underlying positive genetic relationship between yield and N_2_ fixation (%Ndfa) is indistinguishable between older and newer accessions. We note that low-yielding accessions are highly influential to these findings: when the five lowest-yielding pre-1950 accessions are excluded from the comparison, the yield-symbiosis correlations do not differ between the pre- and post-1950 age groups.

Yield in pre-1950 accessions may have been limited by total N to a greater degree than in post-1950 accessions. Greater N-limitation would be consistent with our finding that N_2_ fixation (%Ndfa) is more closely tied to yield increases in yield the pre-1950 vs the post-1950 accessions. Although the efficiency of N_2_ fixation remains an important consideration for pea agriculture (Holmesi et al., 2018), many evaluations of modern varieties do not consider N as limiting for pea yield (Larmure & Munier-Jolain, 2019; Lecomte et al., 2023). While N_2_ fixation (%Ndfa) rate is less consistently associated with yield in more modern pea accessions, we have seen that the slope of the yield-N_2_ fixation correlation has not changed over the 20^th^ century. This means that while breeding seems to have had a stronger effect on yield than on N_2_ fixation (%Ndfa), higher-yielding accessions still tend to get a greater proportion of their N from rhizobia than do lower-yielding accessions. In the MP3 peas, then, there is no evidence of a trade-off between resource allocation to yield or rhizobial mutualists. This is in contrast to the trade-offs that develop between traits in other crops.

We find that yield-seed protein concentration trade-offs are not inevitable in the evolution of mutualisms during crop improvement. A trade-off between seed yield and seed protein concentration occurs in many crops, including quinoa (Curti et al., 2018), brassicas (Peltonen-Sainio et al., 2011), and legumes (Saenz et al., 2023; Skovbjerg et al., 2019). However, in the MP3 peas we examined, this seed protein-yield trade-off is detectable in pre-1950, but not post- 1950 accessions. This trade-off has been detected in peas in some studies (Krajewski et al., 2012; Tar’an et al., 2004), while others have found inconsistent correlations between yield and protein in different populations (Bărbieru, 2021; Crosta et al., 2022; Klein et al., 2020). In the MP3 peas, seed protein concentration is uncorrelated with N_2_ fixation (%Ndfa), and correlates negatively with yield, overall. However, the yield- seed protein concentration trade-off appears to have been broken in the post-1950 accessions, with protein weight increasing along with yield, despite a reduction in seed protein concentration. This suggests such trade-offs are not insurmountable challenges to crop improvement.

In addition to our examination of 20^th^ century breeding, our inclusion of 12 wild accessions in 2020 allows us to examine the impact of pea domestication from its wild progenitor on yield and symbiosis correlations. Theoretical (Porter & Sachs, 2020) and empirical (Martín-Robles et al., 2018) studies posit that wild plants will benefit more from soil mutualists than do their domesticated relatives. Crop plants could evolve a lower benefit from soil mutualists than their wild relatives if N-fertilised soils allow for relaxed selection on the mutualism. Artificial selection in such conditions could favour crop genotypes that allocate resources to reproductive tissues instead of to mutualists. In contrast to this scenario, we observe a trend among wild genotypes whereby the proportion of leaf N derived from rhizobia does not genetically correlate with shoot weight, while in both the pre- and post-1950 domesticated accessions leaf N derived from rhizobia and shoot weight are positively genetically correlated. Furthermore, wild and pre- and post-1950 domesticated peas have indistinguishable numbers of nodules on 6 week old root systems, despite the fact that domesticated pea has substantially greater above ground biomass than wild pea. The fact that domesticated pea accessions are larger and obtain a greater proportion of their N from the same number of nodules, compared to wild pea accessions, is consistent with a scenario in which domestication has increased the efficiency of the mutualism in peas. While we only examined a small sample of wild pea diversity (11 wild accessions) and this limits the conclusions we can draw about the effect of domestication on yield-symbiosis correlations, this finding is consistent with other research that finds improvement to symbiotic N accumulation as a result of domestication in other legumes (Muñoz et al., 2016). The lack of a genetic correlation between yield and N_2_ fixation (%Ndfa) in the wild peas could also reflect a possible lower compatibility between the rhizobia in the agricultural soil we studied with these wild pea accessions than for domesticated peas, as can occur in host-rhizobia pairings (Drew & Ballard, 2010; Kazmierczak et al., 2017; Rigg et al., 2021; Terpolilli et al., 2008) (Carlson et al., 2023).

Several caveats limit our ability to discern the generality of our findings. First, the relationship between yield and symbiotic traits may depend on the population of rhizobia and the soil chemistry at a site, so more broadly distributed studies are required to determine whether the patterns we observe are consistent across locations. For example, incompatible pairings of plant and rhizobia genotypes could lead to an underestimation of the plants’ ability to benefit from the mutualism (Drew & Ballard, 2010; Kazmierczak et al., 2017; Rigg et al., 2021; Terpolilli et al., 2008). Second, the density of rhizobia at a site may influence the outcome of symbiosis. Inoculation can improve the yield of pea crops (Ahmed et al., 2007; Vessey, 2004). However, our field site has not been deliberately inoculated with rhizobia in the last 25 years, so there may have been missed potential for our peas to benefit from inoculation with highly effective rhizobia. Third, soil fertility can impact the benefits of symbiosis. The MP3 peas were N fertilised which can reduce the benefits of the rhizobial mutualism, as it lowers a plant’s demand for fixed N (Regus et al., 2016). While it can be useful to evaluate rhizobial benefit and breed for improved N_2_ fixation in low-N conditions (Rupela & Rao, 2004), some N fertilisation is recommended to maximise pea yields (Pavek & NRCS Pullman Plant Materials Center, 2012). The peas in our experiment were fertilised with an average of 39.2 kg ha^−1^ N across the three years of the study, lower than the recommended 50 kg ha^−1^ (Pavek & NRCS Pullman Plant Materials Center, 2012). Even if legumes aren’t fertilised directly, they are often grown in rotation with crops that are (Pampana et al., 2018).

## Conclusion

Across 87 accessions of domesticated pea, we observe a positive genetic correlation between yield and the proportion of N that plants get from rhizobia. This genetic correlation suggests that selection for yield has enhanced the pea-rhizobium mutualism, rather than disrupted it, as other studies suggest may occur. In fact, the higher-yielding, most agronomically valuable pea accessions also tend to get a larger proportion of their N from the rhizobial mutualism. This positive genetic correlation, coupled with abundant genetic variation in N_2_ fixation (%Ndfa), indicates that there is great potential for simultaneous improvement of both pea symbiotic N_2_ fixation (%Ndfa) and pea yields. Further, since the post-1950 age group exhibits greater yield without the yield-protein concentration trade-off seen in the pre-1950 age group, gains in either yield or protein may not jeopardise the other as they can in other crops. However, a major storm over the course of the three-year experiment damaged plants and resulted in no detected genetic variance for symbiosis traits in 2020 and no genetic correlation between yield and symbiosis traits. As weather extremes become more common, genetic correlations between yield and symbiosis could continue to be disrupted. Extreme weather disturbances could reduce the benefits of symbiosis in agriculture and hamper breeding efforts. Nevertheless, our findings suggest it is possible to breed pea accessions that meet demands for increased yield, seed protein concentration, and nitrogen fixation, and thus reduce agricultural dependence on environmentally detrimental inorganic N fertilizer.

## Author contributions

Niall Millar collected and analysed the data, and wrote the manuscript. Clarice Coyne, Stephanie Porter, and Niall Millar designed and oversaw the experiment. Stephanie Porter edited the manuscript.

## Acknowledgements

We would like to thank Kurt Tetrick, Jennifer Morris, Britton Bourland, and the rest of the USDA team at the Central Ferry Research Farm for the work that went into planting, managing, and harvesting each year of this experiment. We would also like to thank Camille Wendlandt, Christopher Dexheimer, Sherin Kirubakaran, Nathan Crooks, and Ethan Gallegos for their assistance with data collection in 2020. Thanks to Benjamin Harlow and Michael Lott at the Washington State University Stable Isotope Core Laboratory for performing the isotopic analysis of the plant samples. Thanks also to Murray Unkovich and Cathrine Skovbjerg for providing advice on our analysis. Funding was provided by the National Science Foundation (DEB- 1943239 to Stephanie Porter) and the USDA ARS Pulse Crop Health Initiative (to Clarice Coyne).

## Data availability

The data for this study and the code used to analyse it will be archived on Harvard Dataverse.

