## Supplementary Material for "Selection For Yield Enhanced Rhizobial Mutualism In Pea"

**Supplement: Selection for yield is predicted to enhance the rhizobial mutualism in pea**

**Methods supplement**

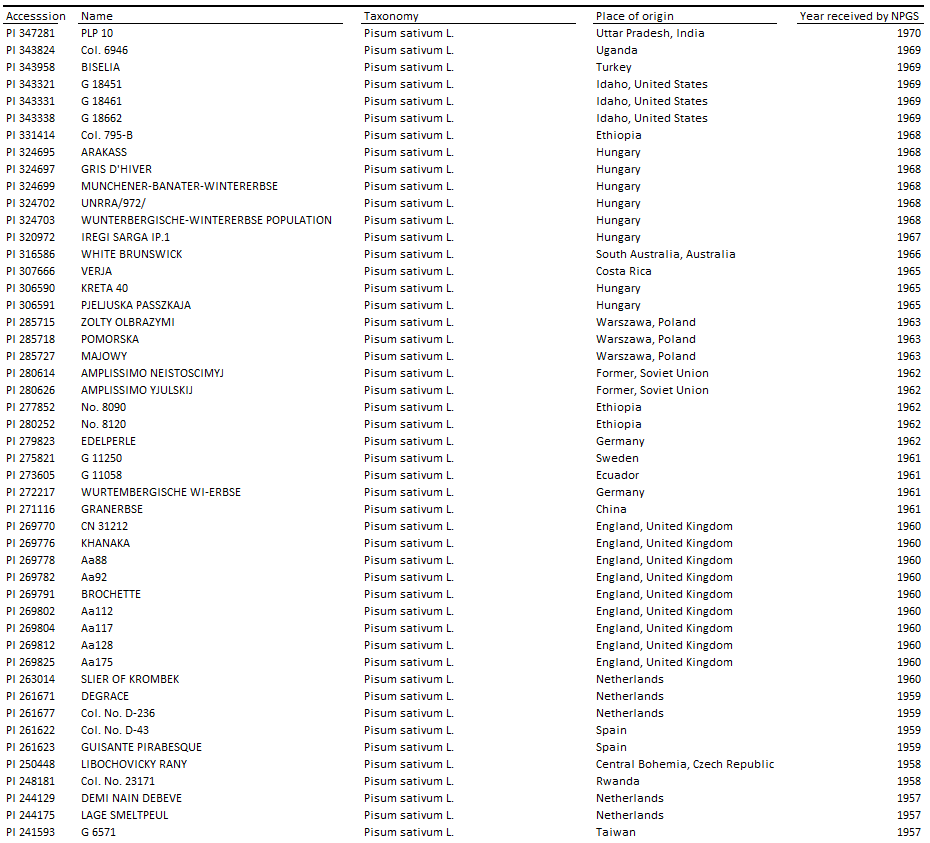

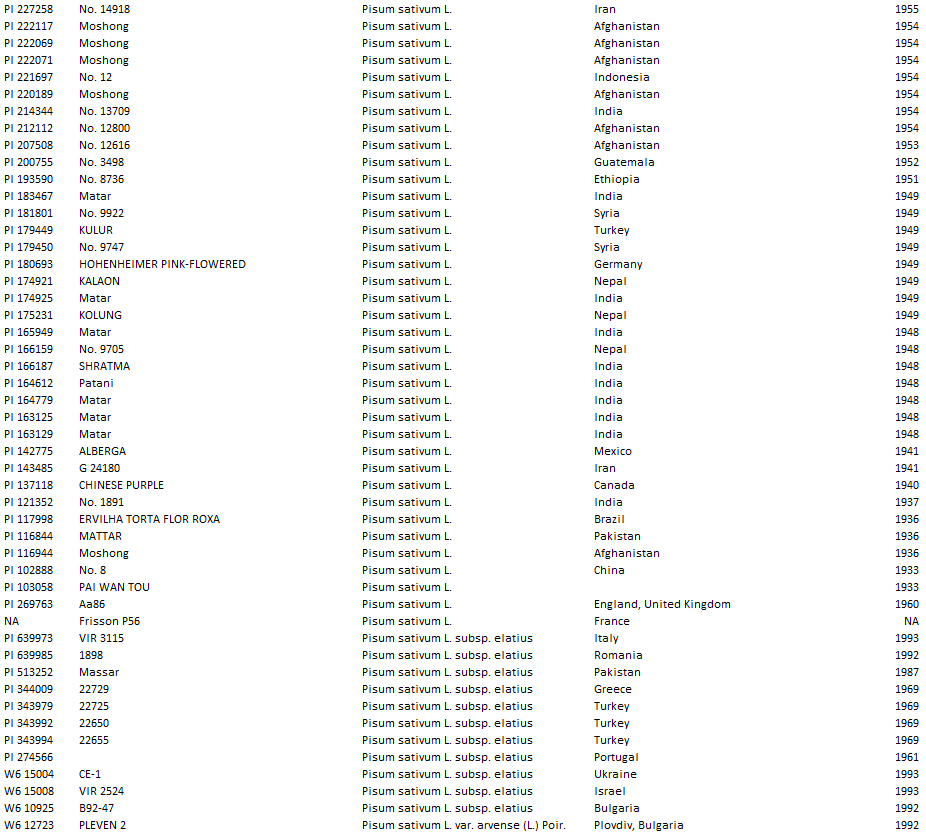

**Table S1: List of accessions used in experiment.** Year received by NPGS refers to the year that the accession was catalogued by the National Plant Germplasm System.

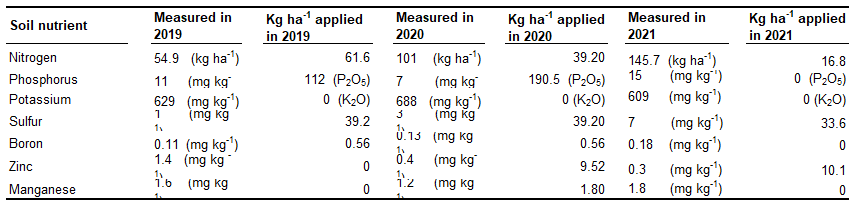
**Table S2: Fertiliser applications by year**

**Results supplement**

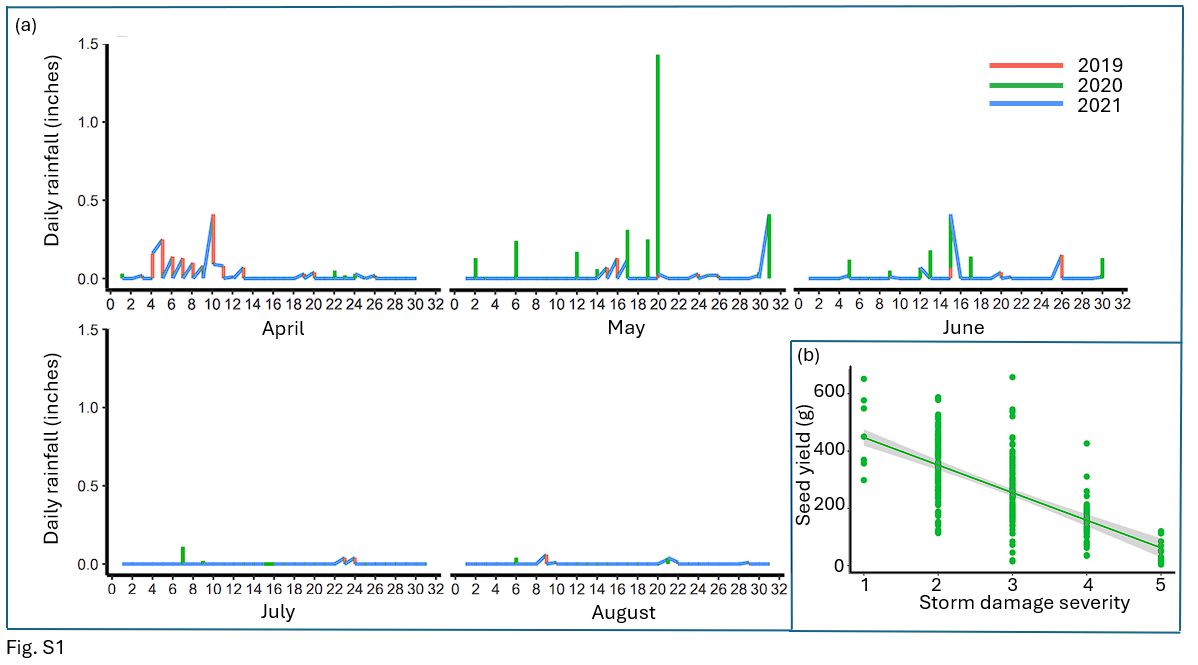

**Fig. S1: May 2020 was an extreme outlier in rainfall due to the rainstorm on the 20^th^. Storm damage severity correlated negatively with seed yield.** Daily rainfall totals per growing season month for 2019, 2020, and 2021 (a), and the correlation between seed yield and storm damage severity (b).

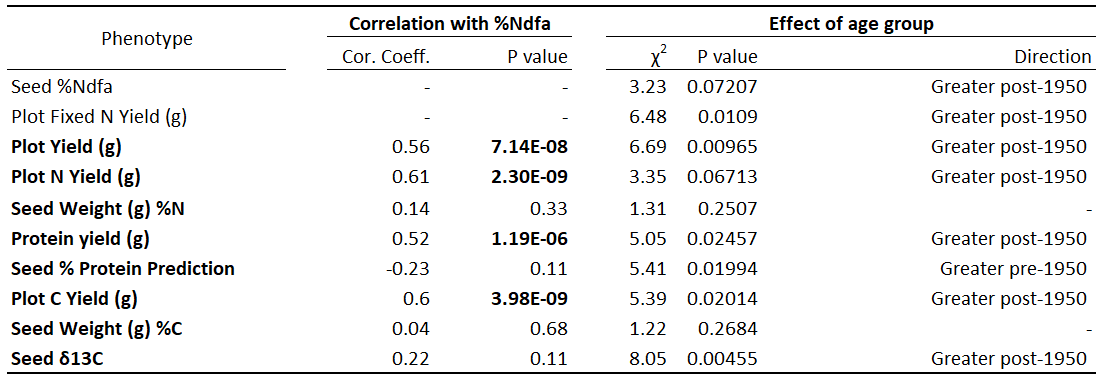

**Table S3: Correlations between seed %Ndfa and yield traits, and the effect of age group on traits, with data from 2019 & 2021.** Pearson coefficients and P values for each %Ndfa-yield trait correlation are shown, as well as the χ^2^ value and P value of likelihood ratio tests of the effect of age group on each trait.

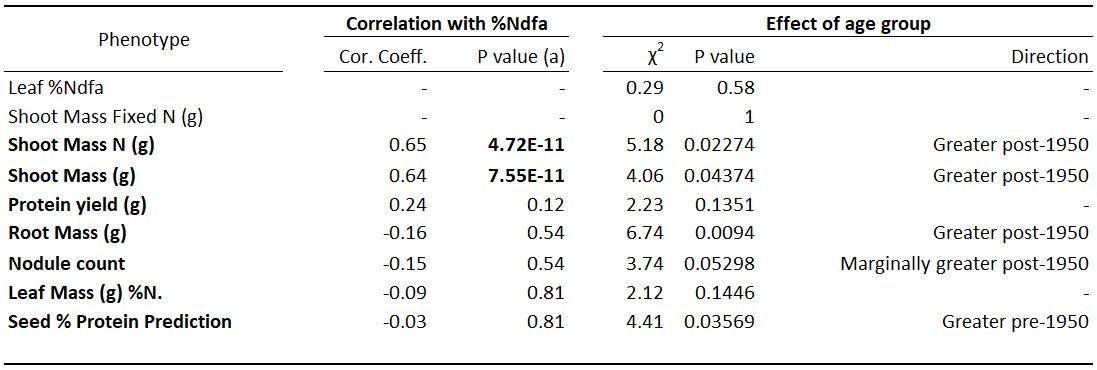

**Table S4: Correlations between seed %Ndfa and yield traits, and the effect of age group on traits, with data from 2020.** Pearson coefficients and P values for each %Ndfa-yield trait correlation are shown, as well as the χ^2^ value and P value of likelihood ratio tests of the effect of age group on each trait.

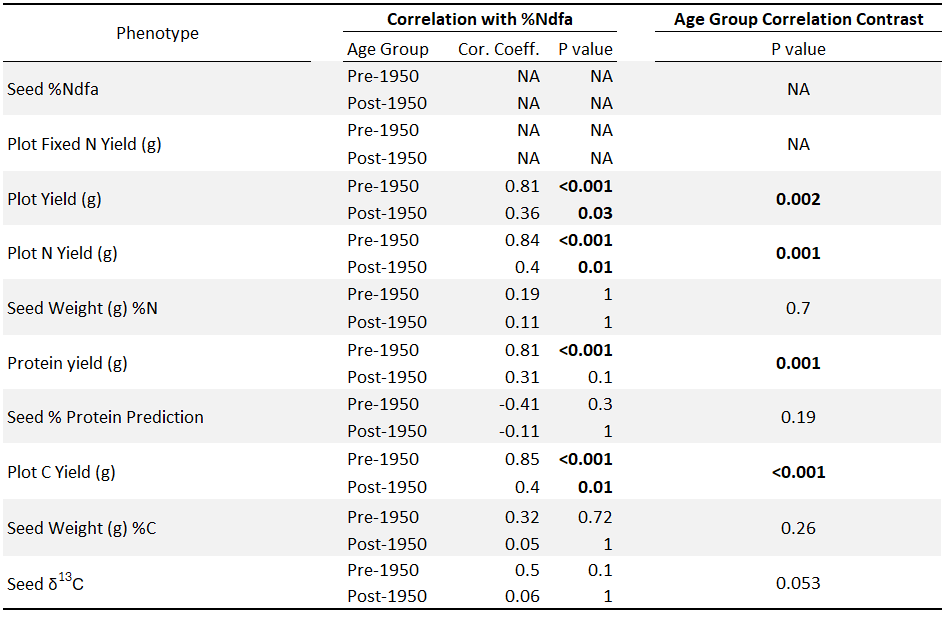

**Table S5: Contrasts between pre- and post-1950 seed %Ndfa- yield trait correlations, with data from 2019 & 2021.** Pearson coefficients and P values of each pre- and post-1950 correlation are shown. The P values of the contrasts between each pre- and post-1950 correlation are also shown.

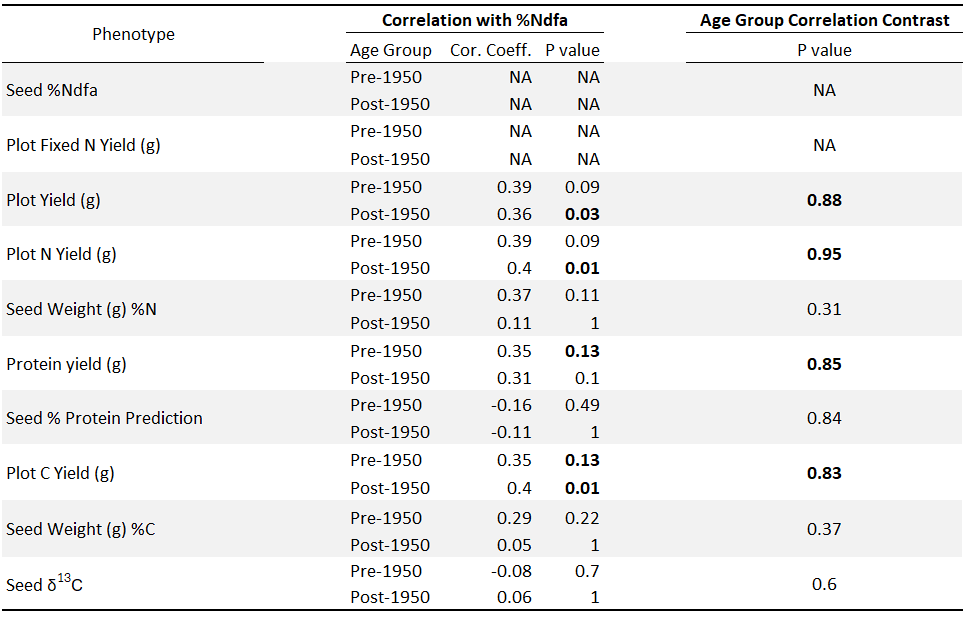

**Table S6: Contrasts between pre- and post-1950 seed %Ndfa- yield trait correlations, with the five lowest-yielding pre-1950 accessions excluded, with data from 2019 & 2021.** Pearson coefficients and P values of each pre- and post-1950 correlation are shown. The P values of the contrasts between each pre- and post-1950 correlation are also shown.

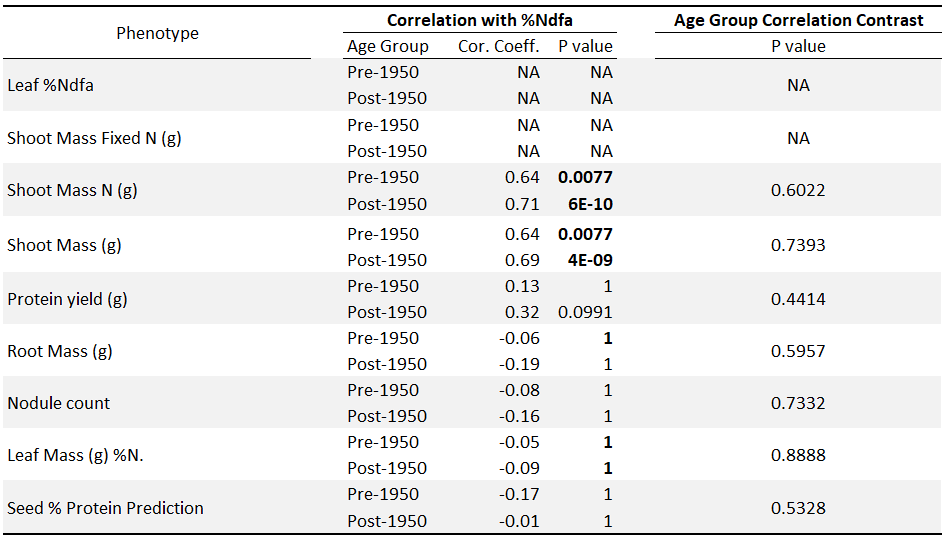

**Table S7: Contrasts between pre- and post-1950 seed %Ndfa- yield trait correlations, with data from 2020.** Pearson coefficients and P values of each pre- and post-1950 correlation are shown. The P values of the contrasts between each pre- and post-1950 correlation are also shown.

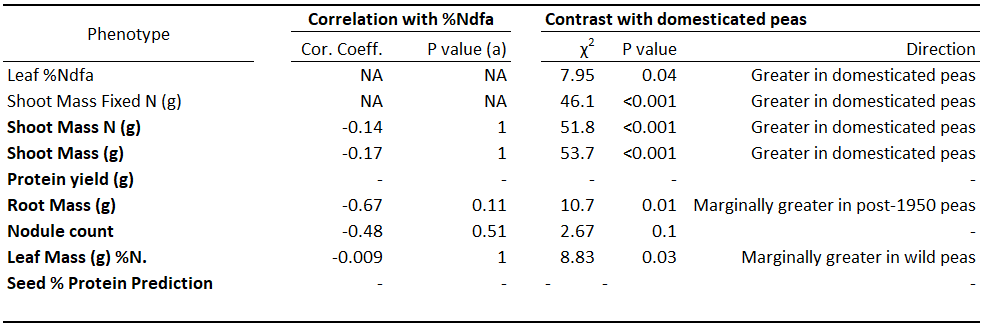

**Table S8: Correlations between leaf %Ndfa and yield traits for wild accessions in 2020, and their contrasts with the domesticated accessions.** Pearson coefficients and P values for each %Ndfa-yield trait correlation are shown, as well as the χ^2^ value and P value of likelihood ratio tests of the effect of domestication status on each trait.

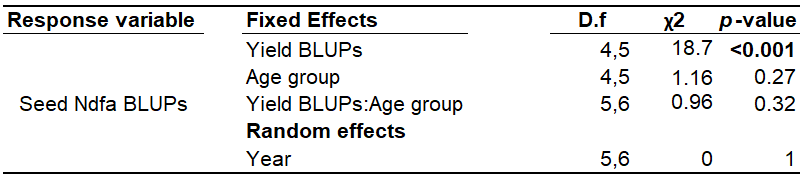

**Table S9: GLMM comparing the slope of the yield-N fixation genetic correlation between pre- and post-1950 accessions.** The significant effect of the yield BLUPs shows the positive genetic correlation between yield and N fixation. The insignificant effect of the yield BLUPs:Age group interaction shows that the slope of the yield-N fixation genetic correlation does not differ between the pre- and post-1950 age groups.
